# Fibroblast Growth Factor 21 (FGF21) creates sugar-specific taste aversion to fructose through action in the brain in mice

**DOI:** 10.1101/2020.01.27.921361

**Authors:** Darko M. Stevanovic, Alex J. Hebert, Bhavna N. Desai, Garima Singhal, Andrew C. Adams, Jeffrey S. Flier, Eleftheria Maratos-Flier

## Abstract

Metabolic diseases such as diabetes and obesity are a growing healthcare concern, and their increasing rates are attributed to increased consumption of carbohydrate-rich diets and sugar-sweetened beverages. Fibroblast growth factor 21 (FGF21) is a complex metabolic regulator, and there is significant evidence that it may play a role in fructose metabolism, driving relative aversion to sweet taste. As such, we examined the relationship between FGF21 and the preferential intake of simple carbohydrates in mice, both as liquid solutions and as dietary additives. Genetic deletion of FGF21 or its obligate co-receptor β-klotho (KLB) had no impact on preference for sugar sweetened solutions. FGF21 overexpression, however, substantially suppressed preference for fructose solutions, but had no effect on glucose or sucrose preference. Infusions of FGF21 also suppressed fructose preference specifically, an effect that was dependent on expression of KLB in the CNS. These results demonstrate that FGF21 creates sugar-specific taste aversion to fructose, which may be mediated by a KLB-dependent pathway in the brain.

**Highlights:**

- FGF21 administration suppresses fructose preference in mice.
- Preference for glucose or sucrose is not affected by FGF21 administration.
- Genetic FGF21 deletion does not enhance fructose, glucose, or sucrose preference.
- FGF21 requires central β-klotho expression to suppress fructose preference.

## 1. Introduction

Fibroblast growth factor 21 (FGF21) is a peptide with complex physiologic actions. It is expressed in many metabolically active tissues, including the liver, pancreas, muscle, and adipose tissue [1–8]. Regulation of expression is also complex, and multiple transcription factors have been implicated in driving changes in responses to dietary challenges, such as fasting, ketogenic diets, lipotoxic diets, protein restriction, and cold exposure [9–15]. Understanding the actions of FGF21 is also complicated due to species differences. For example, FGF21 expression rises dramatically in liver with both fasting and consumption of ketogenic diets (KD) [9,10,16], an effect that is downstream of the transcription factor PPARα. In contrast, in similar diet studies in humans neither fasting nor consumption of KD alter FGF21 levels [17,18].

Intriguingly, genomic analyses in humans associated FGF21 polymorphisms with macronutrient choice. Two synonymous single nucleotide polymorphisms (SNPs) surrounding the FGF21 allele are associated with increases in general carbohydrate consumption [19,20]. This suggests a possible role of FGF21 in carbohydrate metabolism. Such a role is consistent with results from mouse studies demonstrating that chronic consumption of a high fructose diet leads to long-term persistent increases in FGF21 [21,22]. FGF21 is known to play an anti-inflammatory and anti-fibrotic role in the liver, [11,22] and in the absence of FGF21, diets rich in fructose lead to excess hepatic damage [21,23]. Furthermore in humans, while neither ketogenic diets nor fasting alter circulating serum FGF21 levels, acute ingestion of both fructose and alcohol leads to rapid increases in circulating serum levels of FGF21 [21,22,24].

The aggregate data on induction of FGF21 by fructose ingestion and worsened liver pathology in the absence of FGF21 suggested that mice lacking FGF21 might show a relative aversion to fructose. We therefore examined taste preference in mice in the context of genetic FGF21 deletion, genetic FGF21 overexpression, and exogenous FGF21 administration. We were surprised to find that deletion of FGF21 had no effect on preference for sweetened solutions, and that increased preference for sweet was the same in knockout and wild-type mice. Equally surprising was the finding that high levels of FGF21 exclusively suppressed preference for fructose sweetened solutions, but had no effect on glucose, sucrose or saccharin sweetened solutions. Because taste is a complex phenomenon, results with solid diets did not entirely overlap with solution preferences. When mice were presented with diets supplemented with simple sugars, elevated FGF21 levels led to nil consumption of a high fructose diet, relative preference for chow over a high glucose diet, and equalized consumption of chow and a high sucrose diet. However, absolute preference for a high fat, high sucrose diet was observed in all mice, regardless of FGF21 genotype.

## 2. Materials and Methods

### 2.1. Animal Models

The FGF21 total knockout (KO) and the FGF21 overexpressing (OE) mouse lines used in these experiments were generated by Lilly Research Laboratories (Indianapolis, IN). Mice were provided to our group at BIDMC and backcrossed onto the C57BL/6J line (Jackson Research Laboratories, Bar Harbor, ME) at least 10 times before use. Colonies were maintained by breeding heterozygotes, and the mice used throughout the experiments described here were obtained from these breeding pairs. Brain-specific β-klotho knockout mice (BS-KLB KOs) were generated by crossing KLB flox-flox mice (generous gift from Steven Kliewer, Department of Molecular Biology, University of Texas Southwestern Medical Center, Dallas, TX) with the Snap25-CRE line (Jackson Research Laboratories, Bar Harbor, ME). These mice were also backcrossed onto the C57BL/6J line at least 10 times before experimental cohorts were generated by breeding heterozygotes.

Each colony was individually maintained in the Animal Research Facility of the Beth Israel Deaconess Medical Center (Boston, MA), and mice were paired with their wild-type (for FGF21 KO and OE) or their flox-flox (for BS-KLB KO) littermates as controls. For all experiments, mice were housed in a temperature-controlled environment of 24 ± 2 °C under a 12h light: 12h dark cycle (0600-1800h). All procedures were designed in accordance with the National Institute of Health Guidelines for the Care and Use of Animals and were approved by the Institutional Animal Care and Use Committee of the Beth Israel Deaconess Medical Center.

### 2.2. Liquid Diet Preference Assays

Mice were provided with *ad libitum* access to standard lab chow (Formulab Diet 5008 – LabDiet, St. Louis, MO) and two drinking bottles – one containing water and the other containing the experimental sugar solution – for indicated durations. Sweetened solutions were formulated using autoclaved water and either fructose, glucose, sucrose, or saccharin (F0127, G8270, S0389, and 109185 – Sigma-Aldrich, St. Louis, MO). Liquids were contained in chew-proof and drip-resistant drinking bottles (100079433 – Kaytee, Chilton, WI; 61581 – Hagen, Mansfield, MA), and these bottles were mounted in a secure location that prevented them from being tipped over or moved within the cage. Liquid intake was measured daily with a tabletop scale (to the closest 0.1g) and converted to volumetric units for analysis based on the measured density of the individual solutions. The positions of the two bottles within the cage were interchanged daily during liquid intake measurements to prevent a bottle-place preference from confounding sugar preference results.

### 2.3. Peripheral FGF21 Infusions

To prepare the mice for surgery, each mouse was anesthetized with vaporized isoflurane and the surgical area was shaved and cleaned with an alcohol swab and a povidone-iodine solution. A small incision was made on the back of each mouse, and an osmotic minipump (Alzet Model 1002 – Durect Corporation, Cupertino, CA) was implanted in the back subcutaneously. Minipumps were filled with a 4.0 μg/μL solution of human FGF21 (gift of Lilly Research Laboratories), and exhibited a mean pumping rate of 0.25 μL/hour, resulting in a treatment of 24.0 μg of human FGF21 per day for each mouse [25]. Silk sutures were used to close the incision after minipump implantation, and mice were given a 60-hour post-operative recovery period after which they were used in liquid diet preference assays as described above.

### 2.4. Solid Diet Preference Assays

Mice were provided with *ad libitum* access to water and two diets – a control chow diet (Formulab Diet 5008 – LabDiet, St. Louis, MO) and an experimental sugar diet – for indicated durations. Experimental diets consisted of 60% fructose, 60% glucose, 60% sucrose, or 45% fat/17% sucrose (TD.89247, TD.05256, and TD.06685 – Harlan Teklad, Madison, WI; D12451 – Research Diets, New Brunswick, NJ). Diets were provided in weighted jars with lids, and these jars were placed in metal holders that prevented them from being tipped over or moved throughout the cage. Food intake was measured daily using a tabletop scale (to the closest 0.1g) and converted to caloric units for analysis based on the chemical composition of each individual diet provided by the manufacturer. The positions of the two jars within the cage were interchanged daily during food intake measurements to prevent a jar-place preference from confounding sugar preference results.

### 2.5. Statistical Analyses

All reported values are provided as group means + SEM. Statistical significance was determined using a Student’s *t*-test or standard two-way ANOVA, where appropriate, and the resulting p values are indicated in the corresponding figure legends. Analysis tests were performed using Microsoft Excel and GraphPad Prism software, and GraphPad Prism was used to construct all of the included graphs and figures.

## 3. Results

### 3.1. C57BL/6J Mice Exhibit Distinct Sugar Preferences

In order to establish baseline sugar preference levels in mice with normal FGF21 physiology, we conducted preliminary liquid preference assays on experimentally naïve cohorts of C57BL/6J male mice. These wild-type mice (WT) demonstrated a significant preference for fructose solutions over water, even for concentrations as low as 2.5% (Figure 1A). Their total fructose intake increased with the concentration of the solution, and their preference for it over water increased as well. Using a similar experimental paradigm and fresh cohorts of wild-type mice, we then tested preference for glucose (Figure 1B) and sucrose (Figure 1C). In both cases, mice preferred the sugar solutions over water, and this preference increased as the sugar concentration increased. Furthermore, WT mice consume more sucrose and glucose than they do fructose at equivalent concentrations. Although WT mice prefer all three of these sugar solutions over water, their consumption of the solutions does differ based on sugar provided, and these differences suggest that the physiological response to the individual sugars may differ as well.

**Figure 1:**
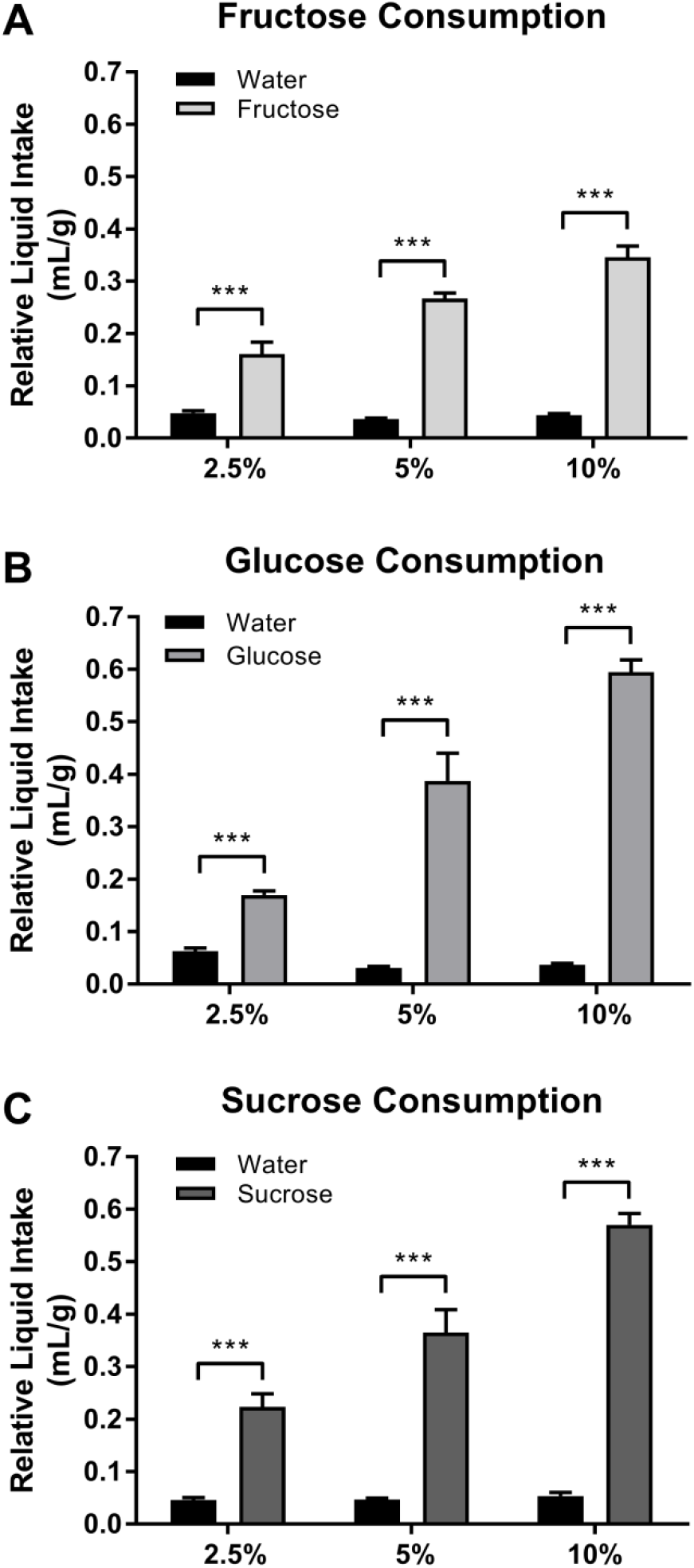
C57BL/6J mice exhibit distinct preferences for sugar solutions. ***A*** Mice significantly prefer fructose solutions of varying concentration (light gray) over water (black). Both their total intake of the fructose solution, and their preference for it over water, increase with the concentration of the solution. ***B-C*** These mice also prefer glucose (*B* – medium gray) and sucrose (*C* – dark gray) solutions over water, and consumption of these solutions increases with concentration as well. This consumption increases at a much sharper rate than it does for fructose, and mice consume more glucose and sucrose than they do fructose for equivalent concentrations. Values represent group means + SEM, with n = 6-10 males for each round of experiments. Significance is indicated by asterisks with *** p < 0.001.

### 3.2. FGF21 overexpression suppresses liquid fructose preference

We next examined the influence of FGF21 on sugar preference using FGF21 OE mice. Mice were provided with a choice between water and a sweetened solution of either 10% fructose, 10% glucose, or 10% sucrose. In contrast to WT mice, which showed a five-fold preference for fructose over water (Figure 2A), FGF21 OEs significantly preferred water, with fructose comprising only 30% of their total liquid intake.

**Figure 2:**
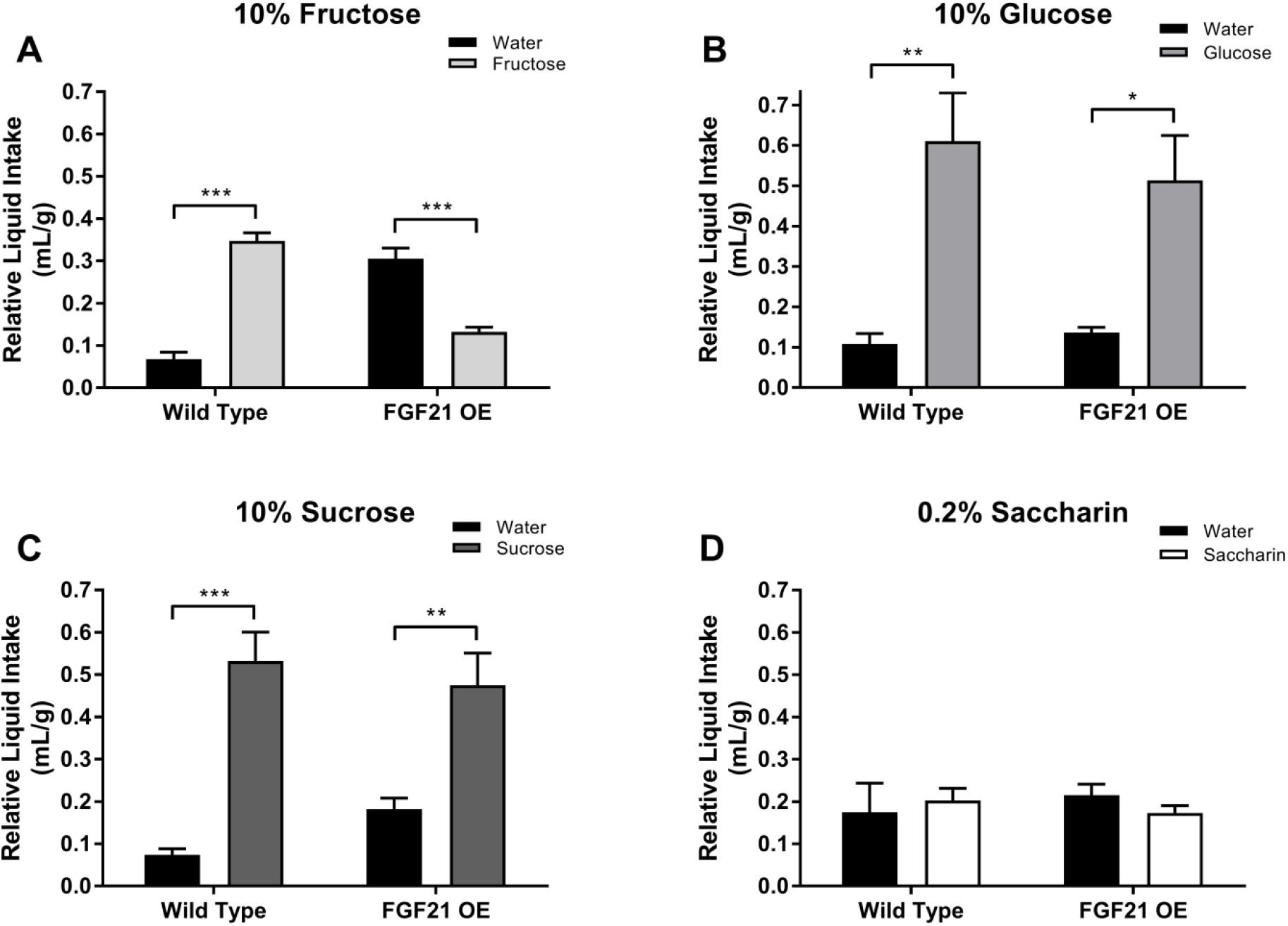
Transgenic overexpression of FGF21 suppresses preference for liquid fructose solutions. ***A*** When given the choice between water (black) and a 10% fructose solution (light gray), WT controls display a significant preference for the fructose solution, while FGF21 OEs prefer water instead. ***B-C*** In contrast, both WT and FGF21 OE mice strongly prefer 10% glucose (*B* – medium gray) and 10% sucrose (*C* – dark gray), consuming significantly more of the sugar solutions than water. ***D*** When provided with a solution of 0.2% saccharin (white), WTs and FGF21 OEs drink equivalent amounts of the artificial sweetener and water, exhibiting no preference for one solution over the other. Values represent group means + SEM, with n = 6 males for each round of experiments. Significance is indicated by asterisks with * p < 0.05, ** p < 0.01, and *** p < 0.001. See Supplementary Figure S1 for daily intake distributions of each solution.

As noted with fructose solutions, WT animals also had strong preferences for the 10% glucose and 10% sucrose solutions, consuming approximately six times as much of the sugar solutions as water (Figures 2B-2C). FGF21 OEs significantly preferred glucose or sucrose over water as well, although the preference ratio was lower (3:1) than their WT counterparts (6:1). To determine if these effects are exclusive to nutritive sweeteners, we also exposed the mice to a 0.2% solution of saccharin. When given the choice between the 0.2% saccharin and water, both WT and FGF21 OE mice alike consume equivalent amounts of the two solutions (Figure 2D).

To examine the effect of FGF21 deletion on a potential analogous increase in fructose preference, we evaluated liquid preference in FGF21 KO mice. A new cohort of WT controls had significant preferences for all four sweetened solutions over water, while mice lacking FGF21 displayed a preference phenotype that was similar to WT animals (Figures 3A-D). Both genotypes consumed twice the amount of glucose and sucrose than they did fructose, and this difference is exacerbated even further to a three-fold change in consumption with saccharin (Figures 3B+3C vs. Figures 3A+3D), which suggests a possible affinity for certain types of sweetness over others. Furthermore, FGF21 KOs and WT controls consume almost identical amounts of liquid throughout these experiments, unlike their FGF21 OE counterparts. For example, both FGF21 KOs and WTs consume 0.6 mL/g of 10% glucose and 10% sucrose, while only consuming 0.05 mL/g of water during each experiment (Figures 3B-3C). This relationship persists for fructose and saccharin, although total liquid consumption for these sweeteners is much lower (0.2-0.3 mL/g for saccharin and fructose vs. 0.6 mL/g for glucose and sucrose). Despite these sugar-specific differences in overall feeding behavior, preference for each of the sweetened solutions over water remains the same between genotypes, indicating that the absence of FGF21 signaling has no effect on mouse preference for liquid sugar solutions.

**Figure 3:**
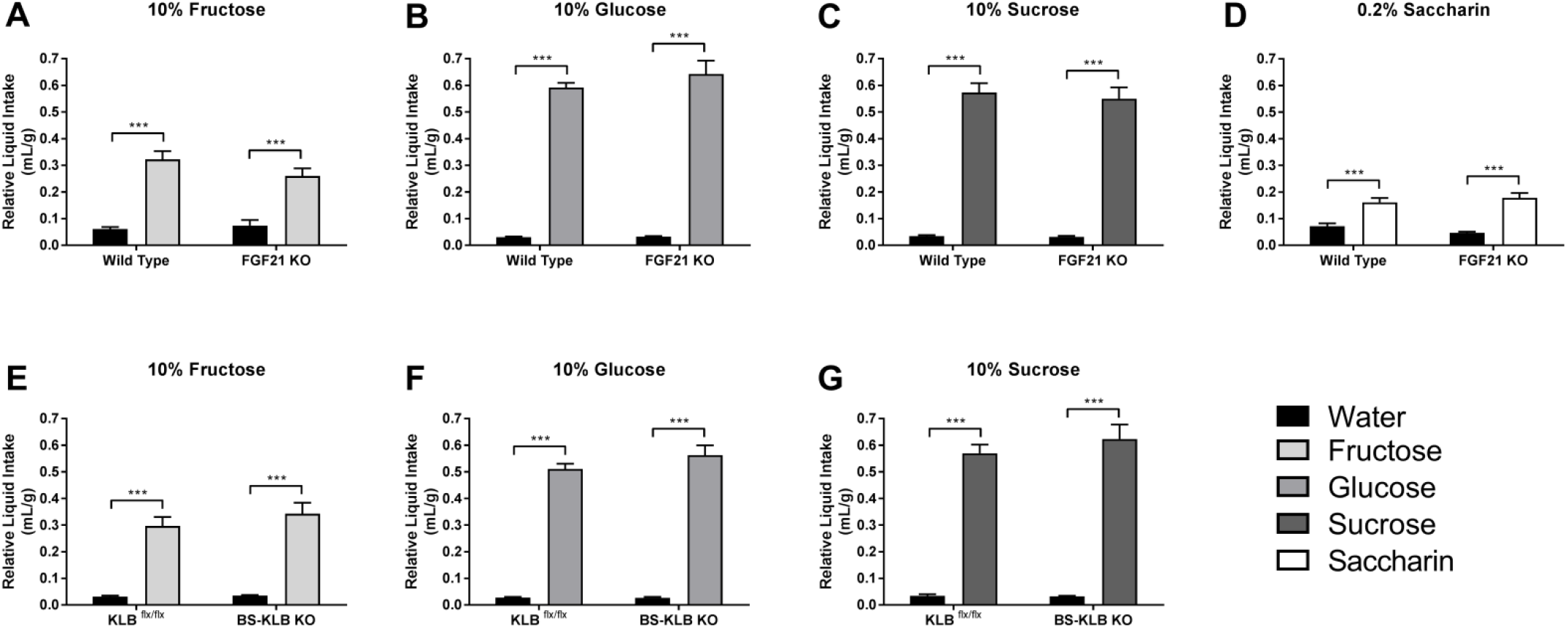
Global FGF21 knockout and brain-specific β-klotho knockout have no effect on liquid sugar preference. ***A-D*** WT mice exhibit a significant preference for 10% fructose *(A* – light gray), 10% glucose (*B* – medium gray), 10% sucrose (*C* – dark gray), and 0.2% saccharin (*D* – white) over water (black), and this phenotype remains unchanged in FGF21 KOs. Additionally, both strains of mice consume larger amounts of glucose and sucrose than they do fructose and saccharin (*B+C* vs. *A+D*). ***E-G*** KLB^flx/flx^ and BS-KLB KO mice also prefer 10% solutions of fructose (*E* – light gray), glucose (*F* – medium gray), and sucrose (*G* – dark gray) over water, and these preferences are nearly identical to those exhibited by FGF21 KOs. Values represent group means + SEM, with n = 6 males for each round of experiments. Significance is indicated by asterisks with *** p < 0.001. See Supplementary Figures S2-S3 for daily intake distributions of each solution.

### 3.3. KLB in the CNS mediates the effect of FGF21

Since taste is a complex phenomenon, mediated by complex neuronal circuitry throughout the CNS [reviewed in 26], we sought to define a potential site of action for the effects of FGF21 on fructose preference. As such, we examined the sugar preferences of mice lacking KLB expression in their CNS (BS-KLB KO). As noted in mice lacking systemic FGF21 expression, BS-KLB KOs had no phenotype when compared to control mice. Both BS-KLB KOs and their KLB^flx/flx^ littermates displayed the same strong preferences for the 10% fructose, 10% glucose, and 10% sucrose solutions over water (Figure 3E-3G). Since the sugar-to-water preferences were similar in FGF21 KOs, BS-KLB KOs, and WT animals, it appears that the absence of FGF21 in otherwise healthy animals has no effect on sweet preference regardless of the sweeter used.

We next evaluated the effect of central KLB expression on the ability of exogenous, pharmacologic FGF21 to suppress sugar preference by treating mice with the hormone via osmotic minipumps. Prior to FGF21 infusions, the mice behaved exactly as they had during the original fructose preference assay (Figure 3E), with both KLB^flx/flx^ controls and BS-KLB KOs exhibiting a strong preference for the 10% fructose solution over water (Figure 4A). At this point, fructose comprised 85-90% of the total liquid intake for each group.

**Figure 4:**
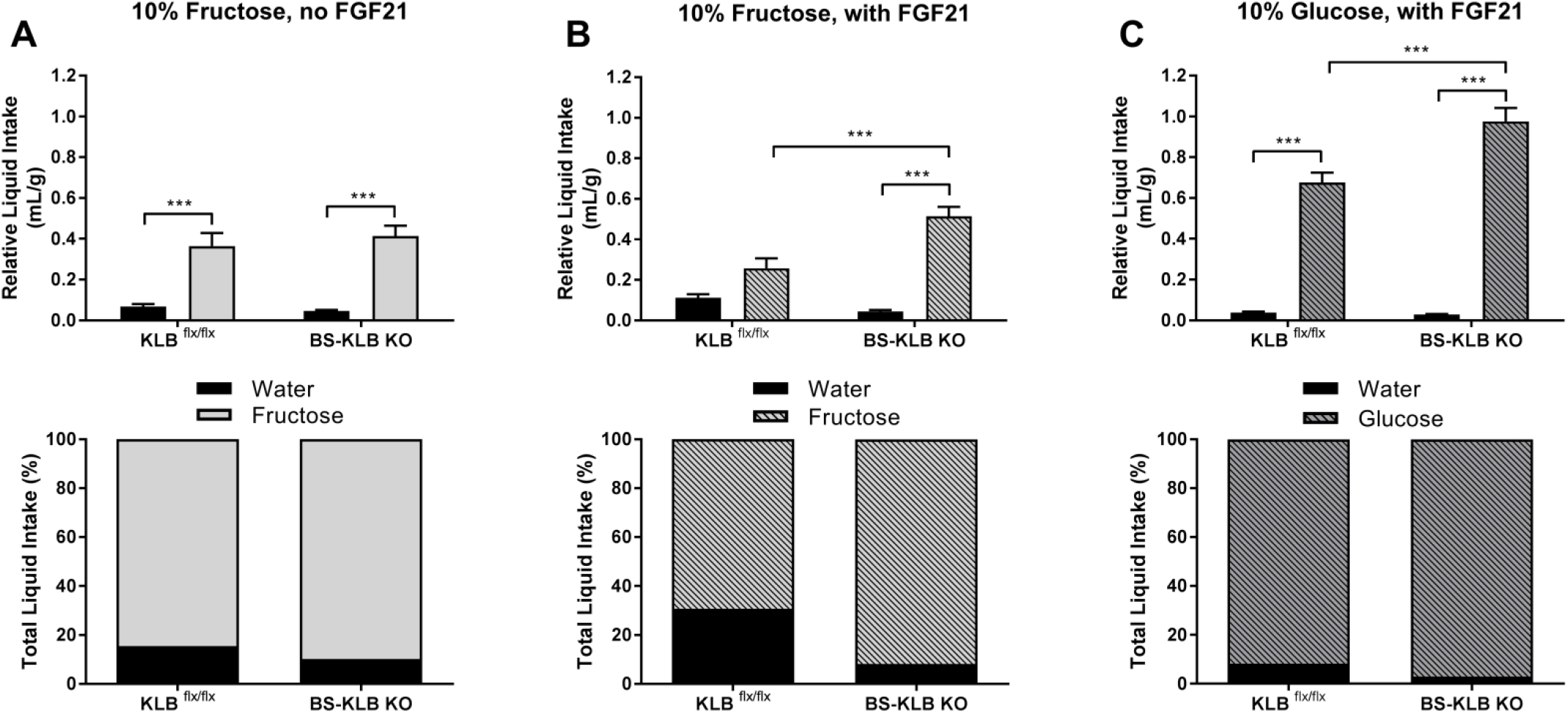
Peripheral FGF21 infusions suppress liquid fructose preference in a brain-specific, KLB-dependent manner. ***A*** In the absence of FGF21 infusions (open bars), both KLB^flx/flx^ and BS-KLB KO mice exhibit similar preferences for 10% fructose (light gray) over water (black). ***B*** Once FGF21 has been infused into the periphery of these mice (hashed bars), the liquid intake of BS-KLB KOs remains unchanged. In contrast, KLB^flx/flx^ controls no longer prefer the fructose solution over water, and they drink significantly less of it than their BS-KLB KO counterparts. ***C*** KLB^flx/flx^ and BS-KLB KO mice both prefer a 10% glucose solution (medium gray) over water when receiving peripheral FGF21 infusions, although the BS-KLB KOs drink significantly more glucose than their control littermates. Values represent group means + SEM, with n = 6 females for each round of experiments. Significance is indicated by asterisks with *** p < 0.001. See Supplementary Figure S4 for daily intake distributions of each solution.

With FGF21 treatment, the liquid intake of the BS-KLB KO mice remained unchanged, with fructose still comprising almost 90% of their total intake (Figure 4B). In contrast, KLB^flx/flx^ mice no longer exhibited a significant preference for the fructose solution over water, and their proportional fructose consumption declined to only 70% of their total liquid intake. These mice drank significantly less fructose than their BS-KLB KO counterparts, and their sugar-to-water preference ratio was five times smaller. Since FGF21 administration can suppress fructose preference in control mice, but not BS-KLB KOs, its effects on fructose preference must be mediated through the brain. Finally, we examined how these FGF21 infusions would impact glucose preference, and we found that both KLB^flx/flx^ controls and BS-KLB KOs alike strongly prefer the 10% glucose solution over water (Figure 4C). While the BS-KLB KO mice did drink significantly more glucose than their control littermates, both cohorts consumed similar amounts of water, and this water comprised a negligible portion of their total liquid intake. The primary effects of FGF21 on sugar preference appear to be fructose-specific, and KLB expression in the brain appears to be a necessary component of this regulatory process.

### 3.4. FGF21 overexpression suppresses fructose preference for solid sugar diets

We then evaluated the effect of FGF21 on preference for sugar sweetened diets, using fresh mouse cohorts. When given the choice between chow and a 60% fructose diet, WT mice appear to be indifferent, consuming equivalent amounts of the two diets (Figure 5A). In contrast, FGF21 OEs ate chow almost exclusively, deriving only 7.5% of their total caloric intake from the high-fructose diet. While indifferent to fructose, WT animals displayed a strong preference for the 60% glucose and 60% sucrose diets, eating twice as much of the sugar enriched diets as they do chow (Figures 5B-5C). FGF21 OEs, on the other hand, preferred chow over glucose with a 3:1 ratio, but consumed similar amounts of chow and sucrose. Finally, we exposed the mice to a diet enriched in both fat and sugar that is commonly used as an obesogenic diet (45% fat, 17% sucrose). Both control mice and FGF21 OEs significantly preferred it, obtaining more than 85% of their daily caloric intake from this 45% fat diet (Figure 5D).

**Figure 5:**
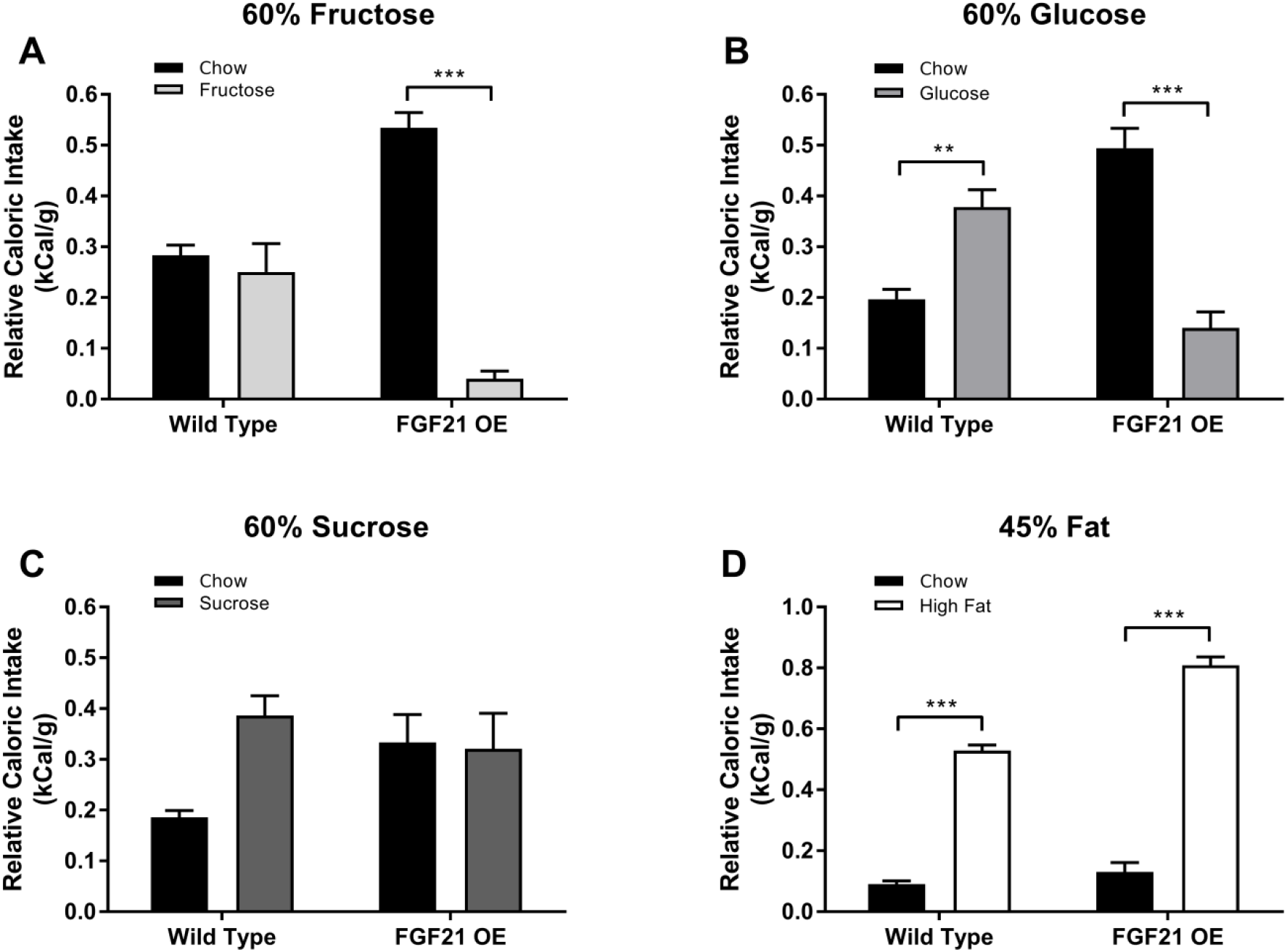
FGF21 overexpression suppresses preference for solid fructose and glucose diets. ***A*** When given the choice between chow (black) and a 60% fructose diet (light gray), WT mice consume equivalent amounts of the two diets, while FGF21 OEs eat chow almost exclusively. ***B*** WTs also prefer the 60% glucose diet (medium gray) over chow, yet FGF21 OE mice display a significant preference for chow instead. ***C*** WT controls consume more of the 60% sucrose diet (dark gray) than chow, but they do not exhibit any significant preference for one diet over the other. FGF21 OEs display no dietary preference for sucrose either, consuming equivalent amounts of the sugar-enriched diet and chow. ***D*** Both WT and FGF21 OE mice strongly prefer a diet composed of 45% fat (white) over chow. Significance is indicated by asterisks with * p < 0.05, ** p < 0.01, and *** p < 0.001. See Supplementary Figure S5 for daily intake distributions of each diet.

Extending our studies to mice lacking FGF21, we repeated these solid preference assays in new cohorts of FGF21 KO mice. WT controls showed a slight, yet insignificant preference for chow over the fructose diet, and FGF21 KOs extended this preference to statistical significance (Figure 6A). Although FGF21 KOs and OEs both exhibited a significant preference for chow over fructose, the extent of this preference was much larger in the FGF21 OEs (Figure 5A vs. Figure 6A). Specifically, FGF21 OEs consumed chow over fructose at a 12:1 ratio, while this preference was reduced to a low 2:1 ratio in FGF21 KOs. WT mice consumed approximately twice as much of the glucose and sucrose diets as they did chow, yet FGF21 KOs ate similar amounts of all three diets (Figures 6B-6C). Preference for these sweetened diets was lost with genetic ablation of FGF21, but we did not see the same degree of active preference suppression that we observed with the FGF21 OEs exposed to liquid and solid fructose diets. We used the 45% fat diet as a control for these experiments as well, and both groups of mice consumed the fat-enriched diet over chow with strong preferences, obtaining more than 85% of their total caloric intake from it (Figure 6D), just like the FGF21 OEs (Figure 5D). Unlike the FGF21 OEs, however, there was no difference in daily or total calorie intake between the two groups of mice.

**Figure 6:**
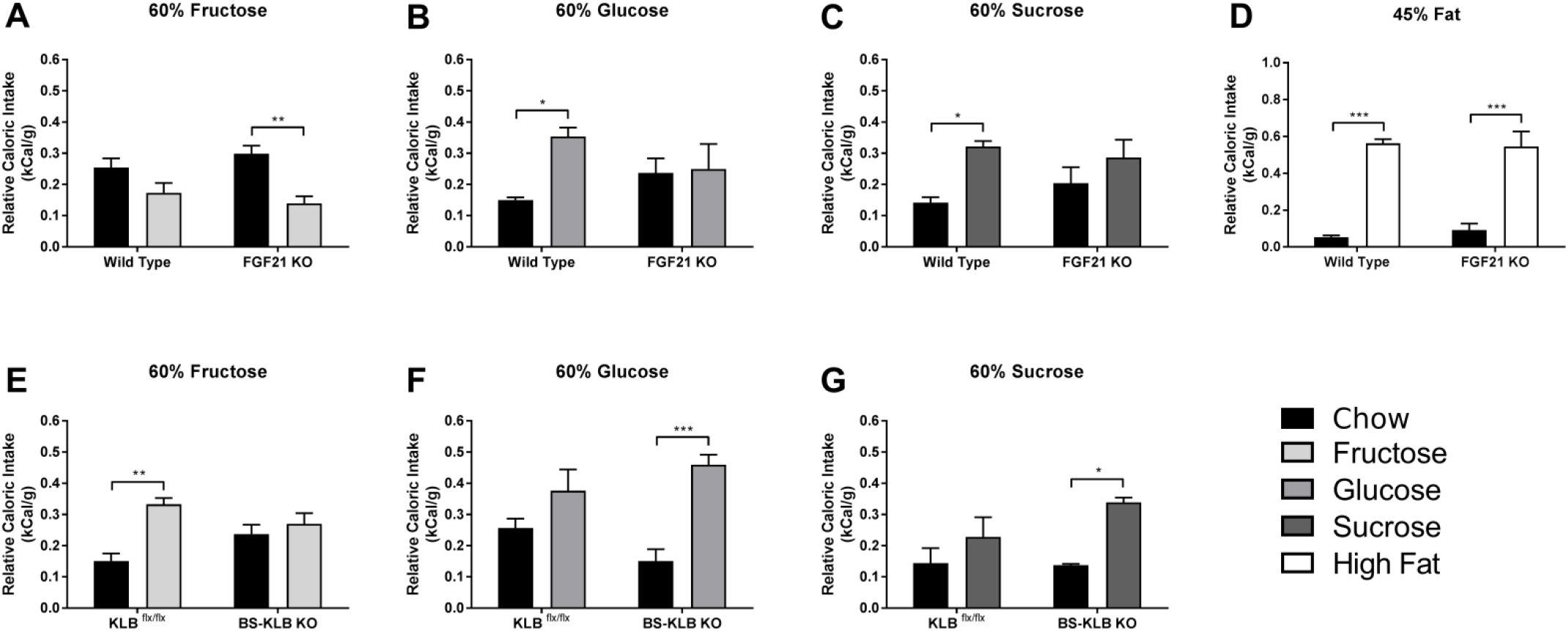
The effects of global FGF21 knockout and brain-specific β-klotho knockout diverge for solid sugar diets. ***A*** FGF21 KOs display a small preference for chow over 60% fructose (light gray), while WT controls do not prefer one diet over the other. ***B-C*** FGF21 KO mice show no preference for either 60% glucose (*B* – medium gray) or 60% sucrose (*C* – dark gray) over chow, unlike their WT counterparts, who prefer the sugary diets. ***D*** Control and FGF21 KO mice alike significantly prefer the 45% fat diet over chow. ***E*** KLB^flx/flx^ mice prefer a 60% fructose diet (light gray) over chow, but this preference is eliminated in BS-KLB KOs, who consume similar amounts of the two diets. ***F-G*** In contrast, BS-KLB KO mice prefer the 60% glucose (*F* – light gray) and 60% sucrose (*G* – dark gray) diets over chow, while their control counterparts display no significant preferences. Values represent group means + SEM, with n = 6 males for each round of experiments. Significance is indicated by asterisks with * p < 0.05, ** p < 0.01, and *** p < 0.001. See Supplementary Figures S6-S7 for daily intake distributions of each diet.

Contrary to our findings with liquid sugar solutions, the solid diet preferences of BS-KLB KO mice diverged from their FGF21 KO counterparts. BS-KLB KOs exhibited no preference for the 60% fructose diet over chow, while KLB^flx/flx^ mice consumed twice as much fructose as they did chow (Figure 6E). BS-KLB KO mice preferred the 60% glucose and 60% sucrose diets over chow, obtaining more than 70% of their total caloric intake from the high sugar diets, unlike their control littermates, who displayed no significant preference for one diet over the other (Figures 6F-6G). While the sugar preferences of BS-KLB KO and FGF21 KO mice do vary when exposed to solid diets, there is still no robust phenotype as the one observed with FGF21 OEs exposed to the fructose diet, suggesting that FGF21 is not responsible for the minor preference fluctuations that occurred throughout these genetic deletion experiments.

## 4. Discussion

FGF21 is regulated by multiple nutritional challenges including fructose which, when administered orally, induces a robust response in both mice and humans [21,24], suggesting that the hormone could play a key role in the metabolism of carbohydrates. Previous reports suggested that FGF21 might also regulate sugar preference and showed that high FGF21 was associated with a relative distaste for sugars, while deletion of FGF21 led to increased sweet preference [27–29].

When we evaluated sweet preference, using both sugar solutions and sweetened solid food, we found significantly more complex responses. When sugars were offered as a liquid, we found that mice lacking FGF21 had the same behavioral preference as that seen in control mice, opting for sugar sweetened solutions when presented with a binary choice of sugar versus water. When we tested sweet preference in mice lacking the co-receptor for FGF21 in the brain, KLB, which precludes FGF21 signaling [30–32], we found that as predicted, these mice had the same phenotype as mice with systemic FGF21 deletion.

Similar to what has been previously reported [27,28], mice overexpressing FGF21 demonstrated a distinct preference for water over fructose. However, we found these mice prefer glucose and sucrose solutions over water, contrary to what has been previously reported. We also found that FGF21 OE mice had equal interest in drinking solutions with the non-nutritive sweetener saccharin compared to water. When we treated control mice with exogenous FGF21, we found that fructose consumption decreased, while this effect was not seen in mice lacking brain KLB receptors.

Taste preference is a complex process that integrates events in the oral cavity, signals from the intestine, and events in the hypothalamus. It is not unexpected that the preference for solid sweetened diets containing either fructose, glucose, or sucrose did not entirely parallel the results with sweetened liquid solutions. While wild-type animals preferred sweetened solutions across the board, the results with solid diets were not as consistent. Wild-type mice did not have a distinct preference for high fructose diets, although they tended to prefer high glucose diets compared to chow. Mice overexpressing FGF21 had significant distaste for high fructose diets, and they preferred chow to high glucose diets as well. Although FGF21 has a small effect on the preferential intake of sugars other than fructose, the effects are dependent on whether it is offered as a solution or as a sweetened diet.

These results are in direct contrast to previous reports that found a preference for carbohydrate-enriched diets in mice lacking FGF21, and also reported that genetic overexpression led to an across-the-board suppression of preference for all sweets [28]. Similar results were reported by another group who found an equivalent decrease in sucrose preference in FGF21 overexpressing mice [27]. In addition, they reported that FGF21 treatment in monkeys also decreased sweet preference, although their experiments were limited to testing preference for an artificial dietary sweetener (saccharin) rather than a dietary sugar. KLB expression is necessary for the regulation of sugar preferences, as reported in all of these studies.

Differences in results may stem from housing in different facilities or subtle differences in the experimental protocol. However, our finding of little to no effect of FGF21 deficiency on taste preference is consistent with previous findings showing little effect of FGF21 deficiency on systemic pathophysiology under baseline conditions of standard chow and housing. Effects are only observed when FGF21 KO mice are challenged by specific pertubations, for example exposure to cold [25]. Pathology can also be induced when mice are challenged with lipodystrophic diets or long term consumption of an obesogenic diet [11–13]. However, in the absence of a specific environmental or dietary challenge, they appear remarkably normal.

Fructose induces FGF21 dramatically in both humans and mice and plays a role in fructose metabolism [21,24]. FGF21 is also induced by sucrose [29], which consisting of fructose and glucose, produces a dimmer response than fructose alone. Thus, it might be expected that in the absence of FGF21, when high fructose has adverse effects on the liver [21], mice could display a relative aversion to fructose and sucrose. However, both our data and data from other groups indicate that aversion is seen consistently and exclusively in FGF21-overexpressing mice [27,28]. While other reports suggest that it is a universal sweet taste aversion, we only observed aversion to fructose. In contrast, FGF21 KO mice, despite the fact that they develop significant pathology when consuming a high fructose diet [21], showed no different preference for drinking fructose compared to WT animals.

While high levels of FGF21 lead to a dislike of fructose solutions, the absence of FGF21 does not have a reciprocal effect. Instead, FGF21’s effects on fructose ingestion appear to be unidirectional in nature with regard to drinking. This is also the case for alcohol consumption, which dramatically induces FGF21 in both mice and humans [22]. While excess FGF21 creates aversion to alcohol, the absence of FGF21 in mice has no effect on consumption.

Our findings suggest that the increase of FGF21 with fructose may be an adaptive metabolic response [21]. In addition to driving metabolism of fructose, the increase in FGF21 may suppress additional ingestion of this sugar. Gustatory sweet preference is directly associated with intake of sweet food [33], so increasing FGF21 in response to consumption may play a role in this preference. This would also explain why the absence of FGF21 has no effect on fructose preference, as an increase over baseline is necessary for a behavioral effect [34,35].

Although our results indicate that FGF21 regulates fructose metabolism by balancing voluntary consumption of the carbohydrate, the exact mechanism it uses to achieve this equilibrium remains unclear. Since FGF21 moderates fructose intake by reducing preference for the sugar, the hormone could influence feeding behavior by altering the short-term desirability of the nutrient. Altering the desirability of a particular stimulus suggests that changes are being made to the neurological reward pathways associated with the stimulus, but FGF21 is not produced in the CNS [36]. Rather, the liver serves as the major organ contributing to circulating levels of FGF21 [2], and is also the major organ to metabolize fructose [37]. Because circulating FGF21 can access the brain, it is plausible that the effects on fructose consumption initiate as yet undefined neurochemical reward pathways. This is consistent with the finding that infusion of FGF21 peripherally only leads to fructose distaste in WT mice, and that this effect is absent when FGF21 cannot signal in the brain, as in BS-KLB KO mice. As such, our results clearly indicate that FGF21 regulates fructose consumption in a neuroendocrine fashion, promoting healthy macronutritional intake of this specific carbohydrate and providing a potential therapeutic target for sugar-related metabolic diseases.

## Acknowledgements

The authors would like to thank Lilly Research Laboratories for providing the FGF21 OE and FGF21 KO mouse lines used throughout these experiments, as well as Steven Kliewer at the University of Texas Southwestern Medical Center for providing the KLB flox-flox mouse line. We are also grateful for the recombinant FGF21 Lilly has provided under a materials transfer agreement.

