## Supplementary Figures for "Fibroblast Growth Factor 21 (FGF21) creates sugar-specific taste aversion to fructose through action in the brain in mice"

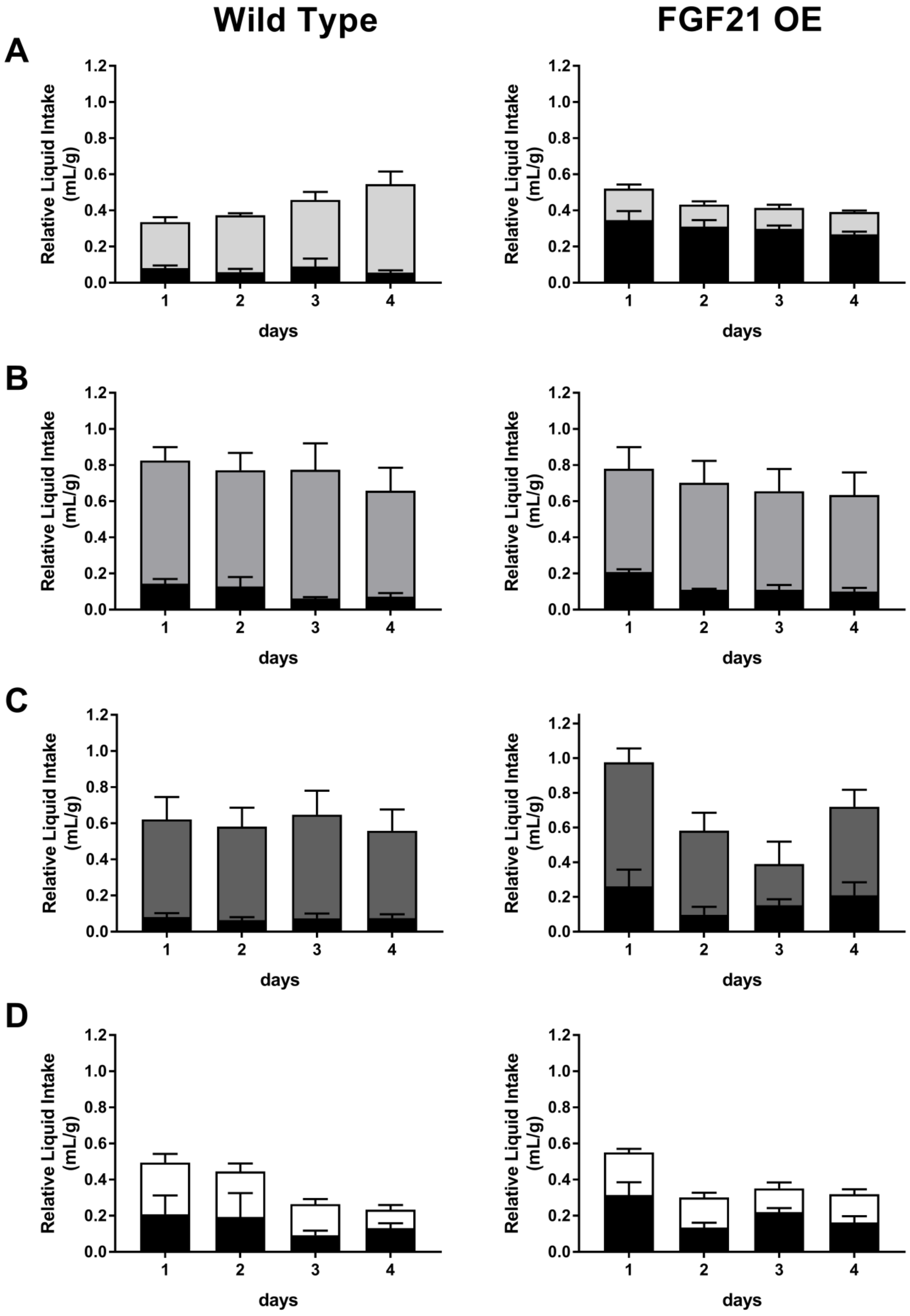


**Supplementary Figure S1: Daily intake distributions for liquid preference assays in FGF21 OEs.** WT (left) and FGF21 OE (right) mice were provided with water (black) and a solution of either 10% fructose (***A*** – light gray), 10% glucose (***B*** – medium gray), 10% sucrose (***C*** – dark gray), or 0.2% saccharin (***D*** – white). Values represent daily consumption group means + SEM, with n = 6 males for each round of experiments.

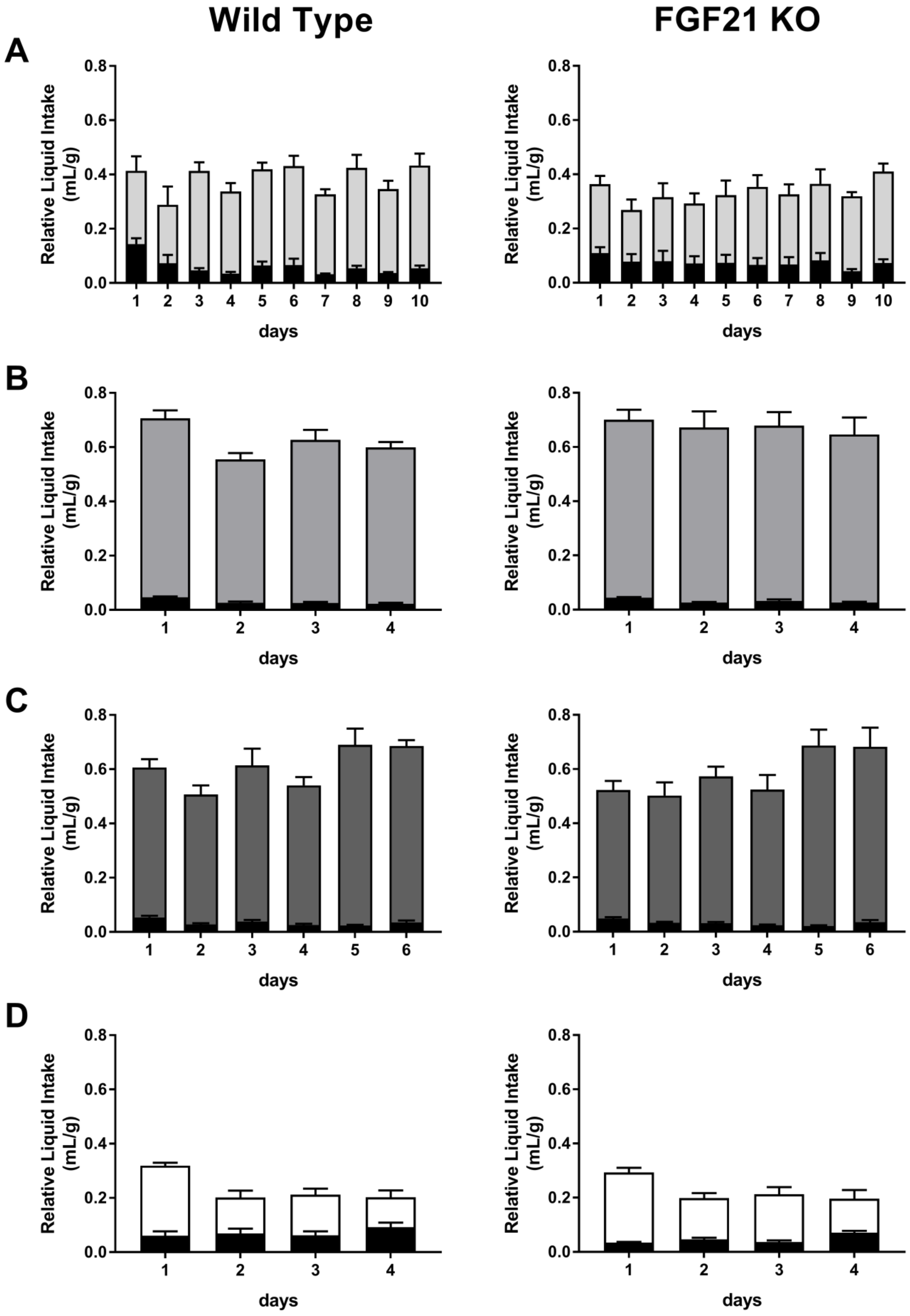


**Supplementary Figure S2: Daily intake distributions for liquid preference assays in FGF21 KOs.** WT (left) and FGF21 KO (right) mice were provided with water (black) and a solution of either 10% fructose (***A*** – light gray), 10% glucose (***B*** – medium gray), 10% sucrose (***C*** – dark gray), or 0.2% saccharin (***D*** – white). Values represent daily consumption group means + SEM, with n = 6 males for each round of experiments.

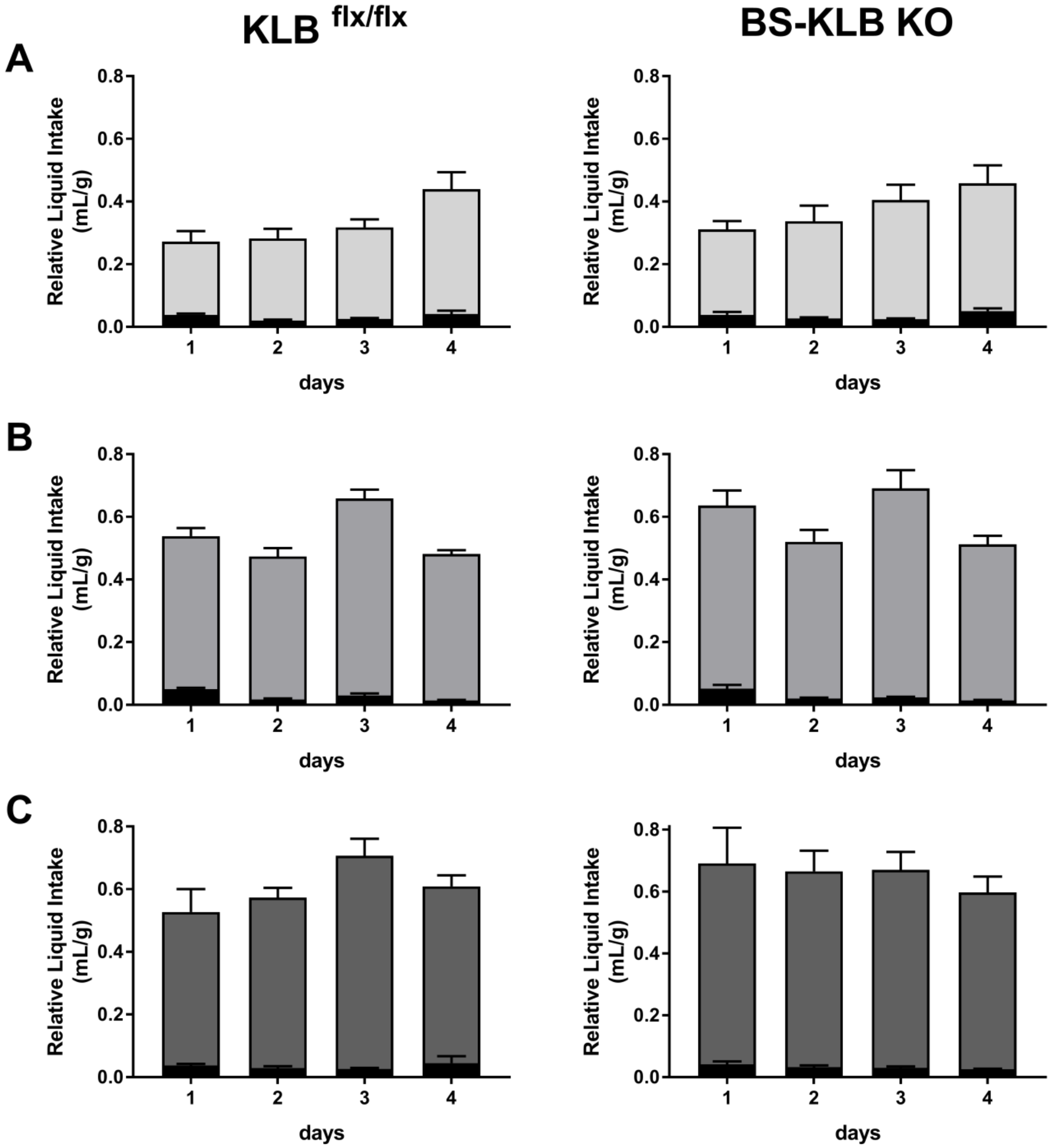


**Supplementary Figure S3: Daily intake distributions for liquid preference assays in BS-KLB KOs.** KLB$^{flx/flx}$ (left) and BS-KLB KO (right) mice were provided with water (black) and a solution of either 10% fructose (***A*** – light gray), 10% glucose (***B*** – medium gray), or 10% sucrose (***C*** – dark gray). Values represent daily consumption group means + SEM, with n = 6 males for each round of experiments.

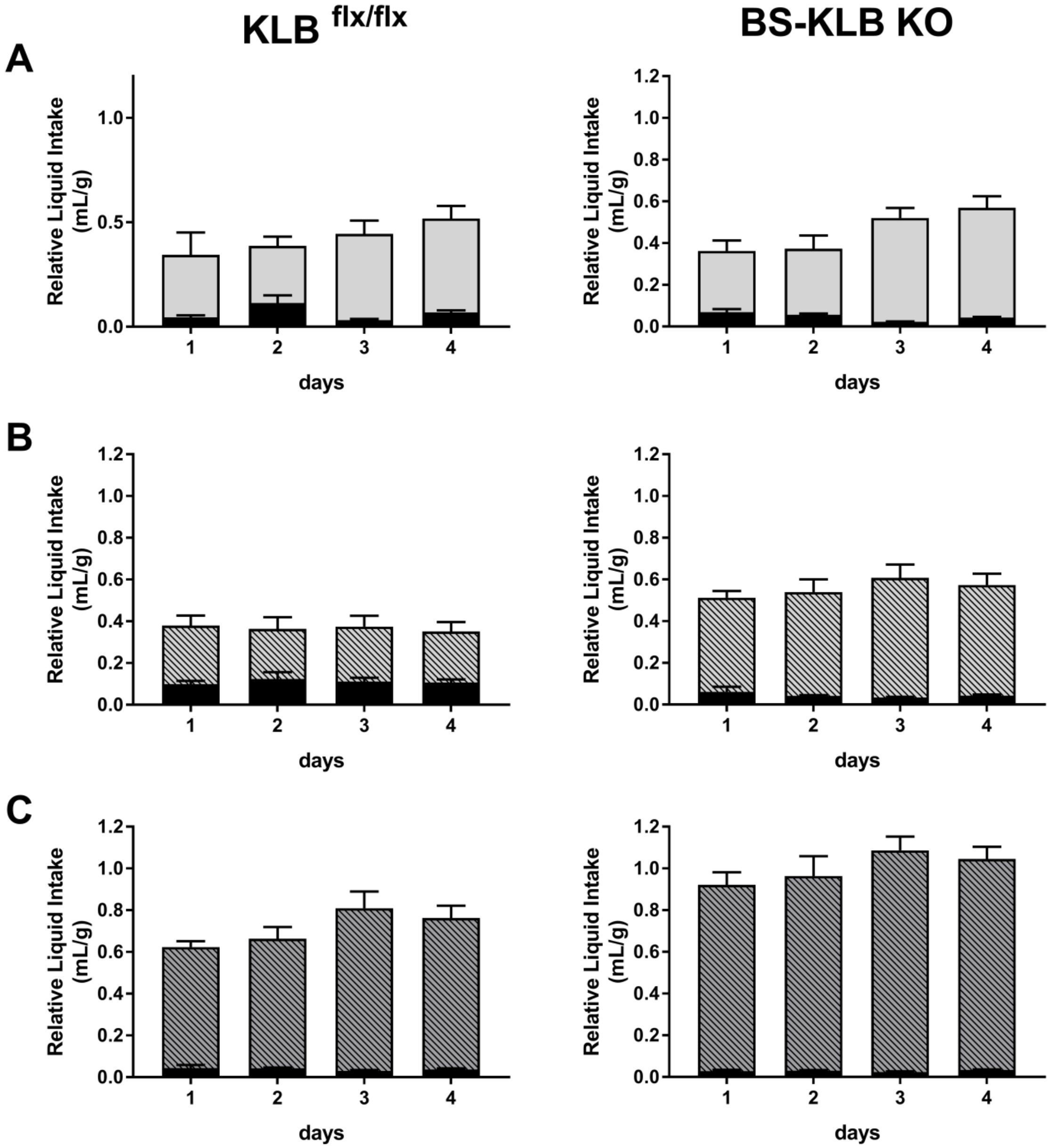


**Supplementary Figure S4: Daily liquid intake for BS-KLB KOs receiving peripheral FGF21 infusions.** KLB$^{flx/flx}$ (left) and BS-KLB KO (right) mice were provided with water (black) and a solution of either 10% fructose (***A,B*** – light gray) or 10% glucose (***C*** – medium gray). Drinking behavior was examined both in the absence (***A*** – open bars) and presence (***B,C*** – hashed bars) of FGF21 infusions, administered subcutaneously via osmotic minipumps. Values represent daily consumption group means + SEM, with n = 6 females for each round of experiments.

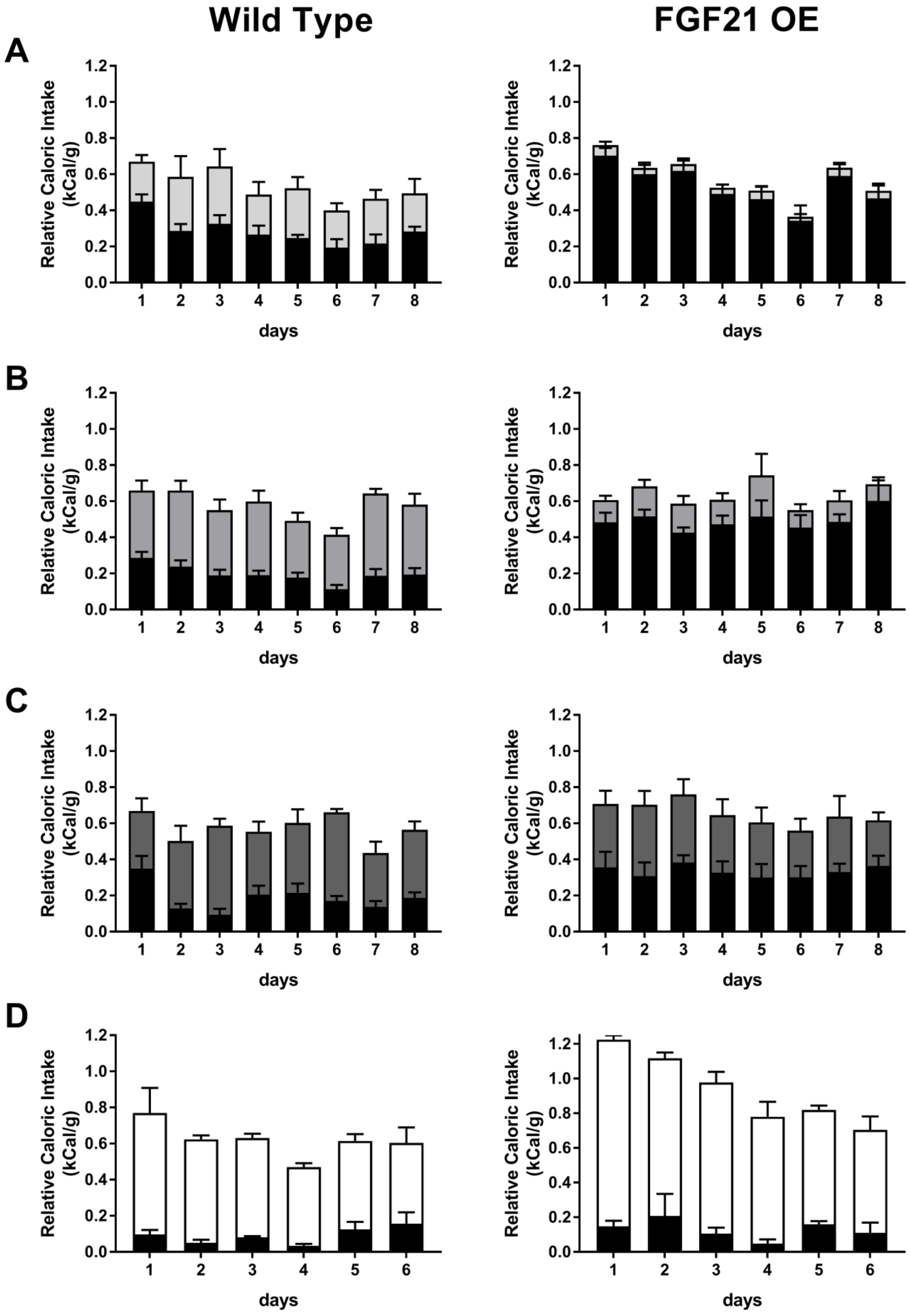


**Supplementary Figure S5: Daily intake distributions for solid sugar diets in FGF21 OEs.** WT (left) and FGF21 OE (right) mice were provided with chow (black) and a diet composed of either 60% fructose (***A*** – light gray), 60% glucose (***B*** – medium gray), 60% sucrose (***C*** – dark gray), or 45% fat (***D*** – white). Values represent daily consumption group means + SEM, with n = 6 males for each round of experiments.

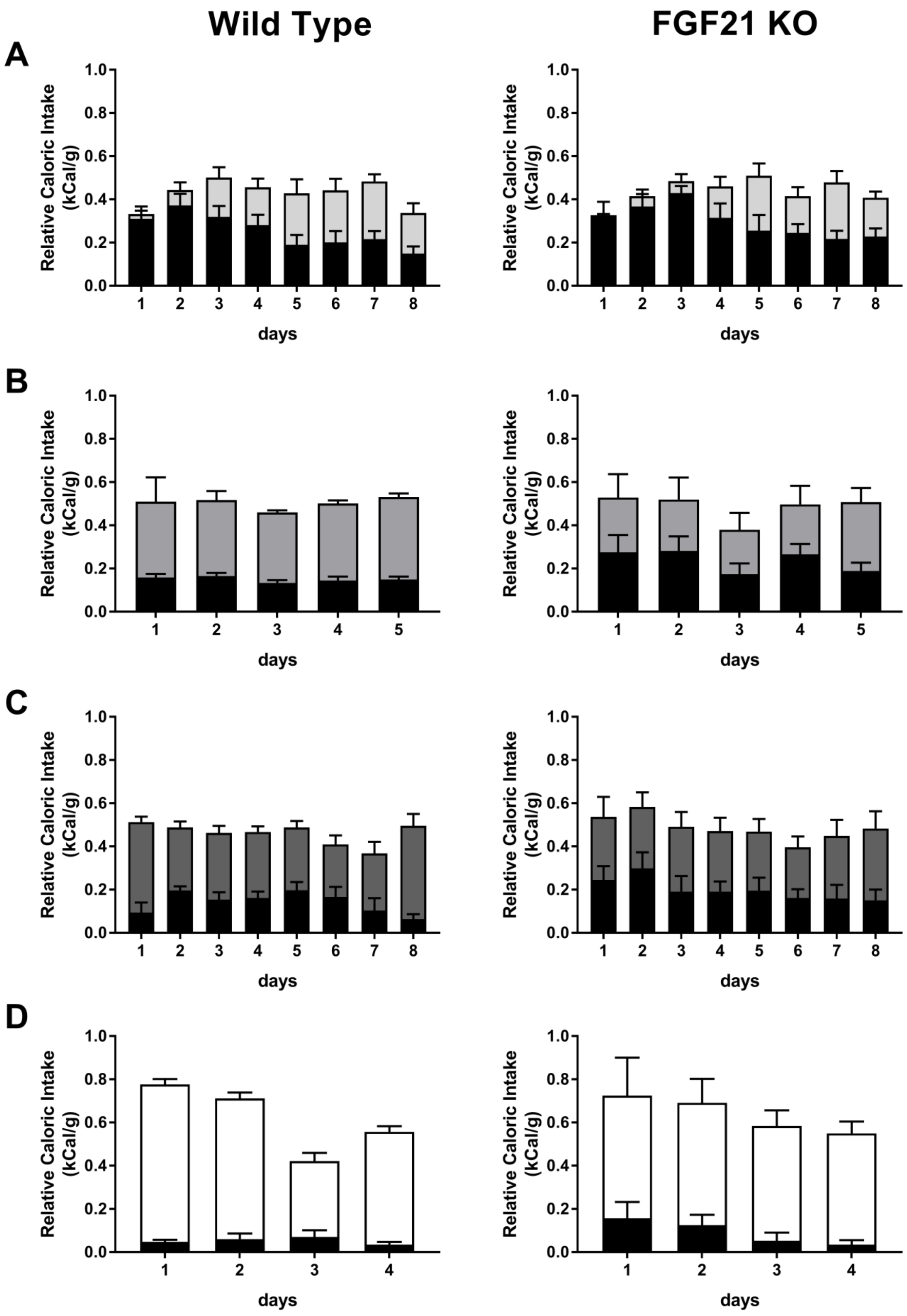


**Supplementary Figure S6: Daily intake distributions for solid sugar diets in FGF21 KOs.** WT (left) and FGF21 KO (right) mice were provided with chow (black) and a diet composed of either 60% fructose (***A*** – light gray), 60% glucose (***B*** – medium gray), 60% sucrose (***C*** – dark gray), or 45% fat (***D*** – white). Values represent daily consumption group means + SEM, with n = 6 males for each round of experiments.

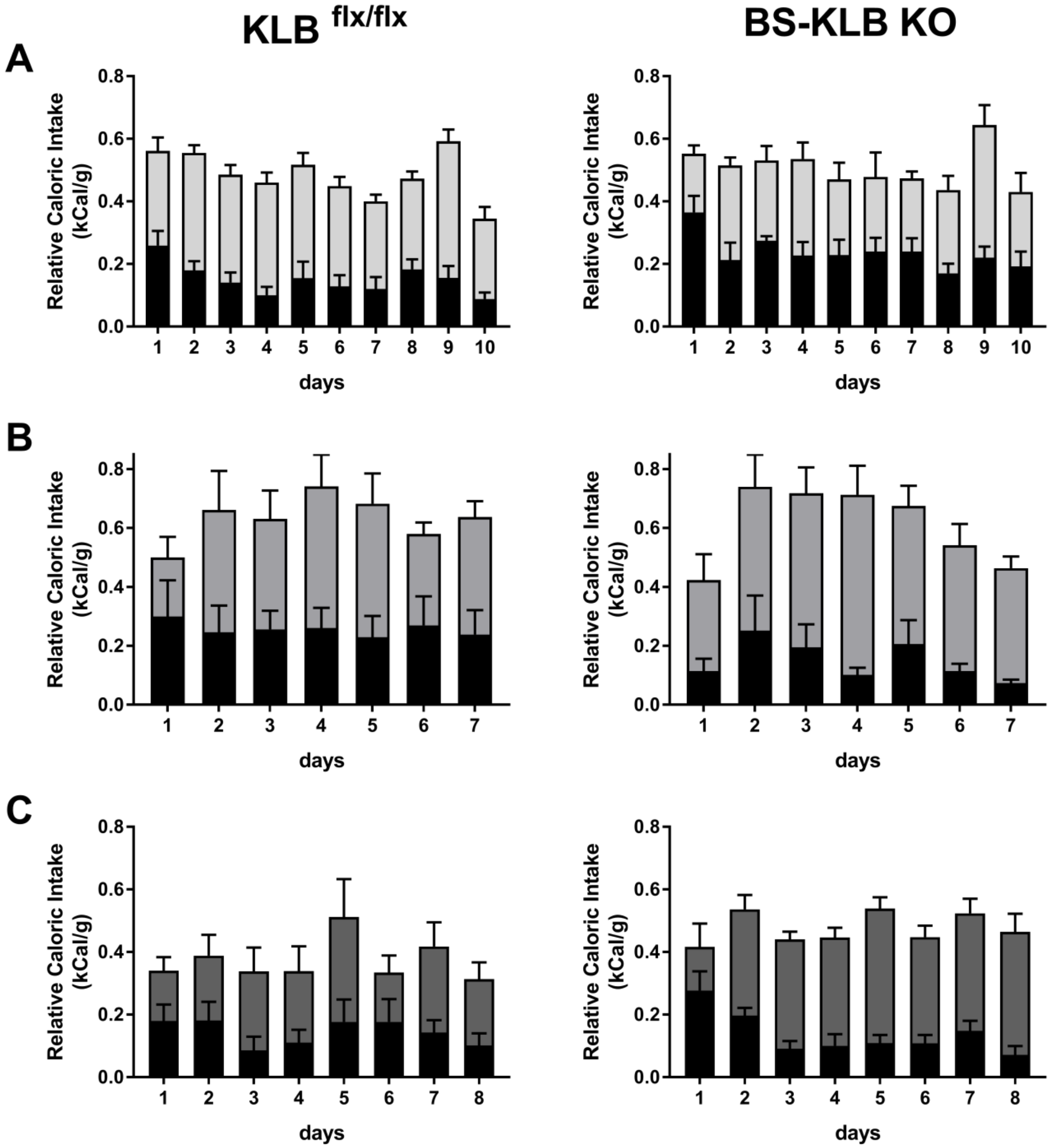


**Supplementary Figure S7: Daily intake distributions for solid sugar diets in BS-KLB KOs.** KLB$^{flx/flx}$ (left) and BS-KLB KO (right) mice were provided with chow (black) and a diet composed of either 60% fructose (***A*** – light gray), 60% glucose (***B*** – medium gray), or 60% sucrose (***C*** – dark gray). Values represent daily consumption group means + SEM, with n = 6 males for each round of experiments.
